# Within and Between-person Correlates of the Temporal Dynamics of Resting EEG Microstates

**DOI:** 10.1101/758078

**Authors:** Anthony P. Zanesco, Brandon G. King, Alea C. Skwara, Clifford D. Saron

## Abstract

Microstates reflect transient brain states resulting from the activity of synchronously active brain networks that predominate in the broadband EEG time series. Despite increasing interest in understanding how the functional organization of the brain varies across individuals, or the extent to which its spatiotemporal dynamics are state dependent, comparatively little research has examined within and between-person correlates of microstate temporal parameters in healthy populations. In the present study, neuroelectric activity recorded during eyes-closed rest and during simple visual fixation was segmented into a time series of transient microstate intervals. It was found that five data-driven microstate configurations explained the preponderance of topographic variance in the EEG time series of the 374 recordings (from 187 participants) included in the study. We observed that the temporal dynamics of microstates varied within individuals to a greater degree than they differed between persons, with within-person factors explaining a large portion of the variance in mean microstate duration and occurrence rate. Nevertheless, several individual differences were found to predict the temporal dynamics of microstates. Of these, age and gender were the most reliable. These findings suggest that not only do the rich temporal dynamics of whole-brain neuronal networks vary considerably within-individuals, but that microstates appear to differentiate persons based on trait individual differences. The current findings suggest that rather than focusing exclusively on between-person differences in microstates as measures of brain function, researchers should turn their attention towards understanding the factors contributing to within-person variation.

---

Neurocognitive networks are believed to enact cognitive functions in real time through dynamic sequences of coordinated brain states (Bressler and Kelso, 2016; Bressler and Tognoli, 2006; Varela et al., 2001). As a consequence, the measurement of the spontaneous activity of global functional brain networks has become a valuable method for studying human cognition and behavior. Contemporary neuroimaging research has increasingly focused on characterizing patterns of spontaneous brain network activity obtained from periods of quiet rest. One important goal of this work is to determine whether intrinsic activity of coordinated brain networks can provide reliable predictors of psychological differences among people in healthy and clinical populations (Dubois and Adolphs, 2016). Aging and disease-related changes in cognitive function are presumed to manifest in structural and functional differences among brain networks (Campbell and Schacter, 2017). Understanding the scale and scope to which intrinsic brain network organization can encode stable, trait-like differences in motivation, affect, and cognition, long argued by personality theorists (e.g. McNaughton and Smillie, 2018), is thus an important topic for continued investigation.

Recent large-sample neuroimaging studies have begun to explore these questions in some detail (Barch et al., 2013). These studies have investigated the contribution of demographic characteristics (Kharabian Masouleh et al., 2019; Smith et al., 2015), personality (Dubois et al., 2018; Nostro et al., 2018), intelligence (Cox et al., 2019), and cognitive function (Lerman-Sinkoff et al., 2017) to patterns of structural and functional connectivity. Much of this work has examined average connectivity patterns of activity derived from whole neuroimaging sessions— so-called “static” functional brain connectivity. Yet recent advances in fMRI-based methodologies suggest that the resting brain dynamically transitions through functional network configurations at much faster temporal scales, on the order of seconds (Abrol et al., 2017; Preti, Bolton, and Van De Ville, 2017). However, even time-varying functional connectivity patterns at this temporal scale are orders of magnitude slower than the millisecond spatiotemporal dynamics accessible through scalp recorded electroencephalography (EEG). As such, EEG may be particularly suited for characterizing the coordinated dynamics of whole-brain neural networks at faster millisecond time scales.

One way to characterize the large-scale dynamics of brain networks is to quantify organized patterns of topographic voltage configurations in broadband scalp-recorded EEG (Michel and Koenig, 2018). When examined across brief timescales, the spatial distributions of certain head-surface voltage topographies can be seen to emerge as quasi-stable patterns in the ongoing signal. During these periods, which typically last between 40 and 120 milliseconds, a particular topographic configuration will tend to predominate, before transitioning rapidly to other momentary quasi-stable configurations. These periods of topographic stability, which have been termed *microstates* (Lehmann et al., 1987; Wackermann et al., 1993), reflect transient brain states of phase-synchronized neuronal activity that can be useful for describing the activity of global cortical networks and their alternating temporal dynamics (for a review of microstate methodology, see Michel and Koenig, 2018).

Any momentary transition in the spatial configuration of the scalp recorded electric field implies, by physical laws, a change in the distribution of active neural generators in the brain (Vaughan, 1982). Because of this, distinct spatial configurations of microstates are taken to reflect the activity of different neuronal networks predominating at that specific moment in time. By extension, the sequencing and temporal dynamics of *changes* in global brain states can be described in terms of the electric field strength and topographic configuration of a succession of microstates over time. Research has consistently identified between 4 and 7 data-driven clusters of microstates that account for a large proportion of the observed variance in EEG time series during periods of quiet rest (Michel and Koenig, 2018). The topographic configurations of clusters of resting EEG microstates are remarkably consistent within and between individuals, and the same clusters have been commonly identified across studies. Moreover, the electrical brain sources of microstates have been suggested to align in part with common fMRI-derived resting state functional networks (Bréchet et al., 2019; Britz et al., 2010; Custo et al., 2017), making microstate analysis a relevant and complementary approach to fMRI techniques for defining the spontaneous organization of the brain.

A rich syntax of neurophysiological parameters can be obtained from the temporal dynamics of microstates at rest, which can be used to characterize ongoing brain network activity. However, only a few studies have examined associations between these microstate temporal parameters and individual differences in age, gender, and psychological dysfunction. To date, measures of the duration, rate of occurrence, and the sequence of specific microstate configurations have been associated with individual differences in age and gender (Koenig et al., 2002; Tomescu et al., 2018), as well as clinical and neurological diagnoses and dysfunction (Rieger et al., 2016; Zoubi et al., 2019). Information regarding the association of microstate parameters with personality attributes or with cognitive performance measures is even more limited. An understanding of the functional significance of microstates, and of the operation of coordinated brain systems more broadly, would benefit from further exploration of what states, traits, or cognitive capacities are associated with microstate temporal dynamics in healthy populations.

In the present report, we assessed a number of within- and between-person correlates of resting EEG microstate temporal dynamics using a large sample of healthy adults. The data for this study were acquired from a publicly available dataset of 227 individuals, comprised of younger and older age-group samples, that was collected between 2013 and 2015 (Babayan et al., 2019). Microstate parameters were derived from 16 minutes of scalp-recorded EEG, collected at rest, from separate eyes closed and eyes open epochs each lasting 8 minutes in duration. Participants were also assessed on a range of demographic, personality, mood state, and cognitive performance measures. We examined associations between these person-level variables and the temporal parameters of microstates, derived from data-driven topographic clustering of voltage maps from the resting EEG data. Specifically, we quantified estimates of (a) the predominance of each microstate configuration in the ongoing EEG signal (i.e., its global explained variance) for each subject; (b) temporal parameters describing the average duration and frequency of each configuration; and (c) the transition probabilities between the occurrence of different configurations in the EEG time series. All participants underwent an extensive medical and psychiatric assessment to screen for active psychological disorders or known health issues.

There were three aims to the present investigation. First, we sought to characterize the sensitivity of microstate parameters and transition dynamics to changes in resting perceptual state by directly contrasting eyes open and eyes closed periods of rest. Second, we examined between-person differences in microstate temporal parameters and transitions as a function of age group (older versus younger adults) and gender. And finally, we explored associations between microstate parameters and measures of personality, mood, and cognitive function included in the larger study assessment battery (Babayan et al., 2019). For personality measures, we selected two commonly used scales that assess motivational tendencies and well-characterized dimensions of personality, respectively: The Behavioral Inhibition and Behavioral Activation scale of Carver and White (1994) and the NEO Five Factor Inventory of Costa and McCrae (1992). In addition, mood state was assessed using the Multidimensional Mood State Questionnaire (Steyer et al., 1997); and attentional performance was indexed with three modules of the Test of Attentional Performance (Zimmermann and Fimm, 2002), designed to assess cognitive facets of psychomotor alertness, stimulus-response compatibility, and working memory.

It is well established that the spatiotemporal dynamics of the brain can vary considerably within as well as between individuals (Van Horn et al., 2008). Such rapid changes in cortical network dynamics can be presumed to subserve momentary shifts in perception and ongoing cognitive function. By a similar logic, the temporal dynamics of microstates might best reflect within-person variability in internal states and cognitive processes that manifest over fast temporal scales. To address this question, we quantified the degree to which temporal parameters of microstates would vary *within* as opposed to *between* individuals. We also examined how microstate parameters were influenced by an individual’s resting perceptual state (eyes closed vs. eyes open), or differed as a function of specific topographic microstate configurations.

Reliable age-related differences have been observed across a number of neuroimaging modalities (Campbell and Schacter, 2017). On the basis of these and other studies reporting age-related differences in microstates (i.e., Koenig et al., 2002; Tomescu et al., 2018), we also expected to find substantial age group differences in microstate dynamics. However, given the exploratory goals of the present study, we had no specific hypotheses regarding which measures of microstate temporal dynamics or microstate configurations would be associated with specific individual differences in gender, personality, mood, or attentional performance. It is plausible that properties of brain systems identifiable through microstates might encode stable trait-like differences in at least some of these measures. Along these lines, it is worth noting a recent study by Liégeois and colleagues (2019) that contrasted static and dynamic fMRI connectivity patterns. These authors reported that time-varying patterns of network connectivity were more strongly predictive of cognitive task performance than were static measures of connectivity (Liégeois et al., 2019), but that static and dynamic functional connectivity were more or less equally predictive of self-report outcomes of affect and life satisfaction. Therefore, we tentatively predicted that measures of cognitive function might more reliably correlate with microstate dynamics than would self-report measures of personality or mood.

## Methods

### Participants

Two hundred twenty-seven participants were recruited as part of the Max Planck Institute Leipzig Mind-Brain-Body study (Babayan et al., 2019). Recruitment targeted two age groups: younger adults between 20 and 35 years old (*N* = 153, 45 females, mean age = 25.1 years, *SD* = 3.1) and older adults between 59 and 77 years old (*N* = 74, 37 females, mean age = 67.6 years, *SD* = 4.7). All participants underwent an extensive medical and psychological screening procedure prior to study inclusion, and were tested at the Day Clinic for Cognitive Neurology of the University Clinic Leipzig and the Max Planck Institute for Human and Cognitive and Brain Sciences in Leipzig, Germany. Further details regarding participant recruitment and eligibility are reported by Babayan and colleagues (2019). All participants provided written informed consent prior to study participation, received monetary compensation for their time in the study, and agreed to their have data shared anonymously. Data were collected and shared by Babayan and colleagues (2019) in accordance with the Declaration of Helsinki and the study protocol was approved by the ethics committee of the University of Leipzig (reference #154/13-ff).

### Procedure

Study assessments took place over two days, with each session lasting approximately 4 hours (see Babayan et al., 2019, for full assessment and data collection details). The first assessment (day 1) included administration of a battery of cognitive tasks, followed by MRI scanning and collection of blood pressure, anthropometric measurements, and blood samples. The second assessment (day 2) included a mood state questionnaire, the acquisition of 16 minutes of resting EEG, and administration of psychological questionnaires and instruments, as well as a psychiatric interview (SCID; Wittchen et al., 1997). Participants were also invited to participate in a follow-up assessment at a later date. Resting EEG was recorded from 216 participants in a sound attenuated chamber. Each recording was divided into 16 contiguous 1-minute blocks, with alternating eyes closed and eyes open conditions, beginning with the eyes closed condition. Participants were seated in front of a computer screen and, for each block, were instructed to remain awake and to sit still with their eyes closed, or to sit still with their eyes open while fixating on a black cross presented on a white background. Presentation software (Neurobehavioral Systems Inc., USA) was used to notify participants of changes between blocks.

### EEG Data Collection and Processing

EEG was recorded from a 62-channel active electrode cap (ActiCAP, Brain Products GmbH, Germany), with 61 channels in the international 10–10 system arrangement. One additional electrode was placed below the right eye to record vertical eye movements. The reference electrode was located at electrode position FCz and the ground was located at the sternum. Data were acquired with a BrainAmp MR plus amplifier (Brain Products GmbH, Germany) at an amplitude resolution of 0.1 µV and sampling rate of 2500 Hz, and were bandpass filtered online between 0.015 Hz and 1k Hz. Following acquisition, the EEG were downsampled offline to 250 Hz and bandpass filtered between 1 and 45 Hz with an 8th order Butterworth filter. Eyes closed and eyes open epochs were concatenated separately, creating one 8-minute segments per condition.

Preprocessed EEG recordings from the Mind-Brain-Body study were made available for use by interested researchers on a data-sharing repository (https://ftp.gwdg.de/pub/misc/MPI-Leipzig_Mind-Brain-Body-LEMON/). Portions of these preprocessed data have already been used in at least two investigations of EEG oscillatory dynamics, by Mahjoory et al., 2019, and by Schaworonkow and Nikulin, 2019. As reported by Babayan and colleagues (2019), outlier channels with poor signal quality, extreme peak-to-peak deflections, or large bursts of high frequency activity were excluded based on visual inspection. Next, principal component analysis (PCA) was used to reduce the dimensionality of the data by retaining components that explain 95% of the total data variance. Infomax independent component analysis (ICA) was then used to remove components reflecting eye movements, eye blinks, or heartbeat-related signal contaminants, and the remaining independent components (*M* = 21.4 components, range: 14–28) were reconstructed. Thirteen participants were excluded due to missing event information, errors in sampling rate, or insufficient data quality, leaving 203 participants with usable EEG.

We gathered the 406 preprocessed EEG recordings for these 203 participants from the data-sharing repository (i.e., one recording for each eyes closed and each eyes open condition). We then interpolated missing electrodes based on spherical spline interpolation to fit a 64-channel montage and re-referenced to the average using the Cartool software toolbox version 3.7 (Brunet et al., 2011). To constrain our analyses to healthy individuals, we excluded an additional 12 participants because the psychiatric interview identified possible untreated psychological diagnoses (e.g., substance abuse or unspecified hallucinations). This left 191 participants for use in the current analysis, with an average of 7.83 minutes (*SD* = 0.51) of eyes closed and 7.77 minutes (*SD* = 0.53) of eyes open resting EEG.

### Topographic Segmentation and Microstate Parameter Estimation

We conducted topographic segmentation of each of the 382 individual EEG recordings (191 eyes closed, 191 eyes open) to identify periods of quasi-stable scalp voltage configurations. This was achieved through an adapted *k*-means clustering method, implemented in Cartool (Brunet et al., 2011), that determines the optimal number of clusters (*k*) that can account for the greatest global explained variance (GEV) in the time series while using the smallest number of representative topographical maps (Michel et al., 2009; Murray et al., 2008). First, topographic voltage maps were generated at local maxima (peaks) in the global field power (GFP) time series. This was done separately for each individual recording. GFP is a reference-independent measure of voltage potential (µV) that quantifies the strength of the scalp electric field at a given sample of the recording, equivalent to the standard deviation of amplitude across the entire average-referenced electrode montage (Skrandies, 1990). GFP peaks were used to generate the initial maps for clustering so as to maximize topographic signal-to-noise ratio. Because GFP peaks tend to reflect moments of high global neuronal synchronization, they provide optimal representations of the momentary quasi-stable voltage topography (Koenig and Brandeis, 2016).

#### Clustering of voltage maps

Iterations of *k*-means clustering proceeded as follows for each recording. For a given participant and condition (eyes closed or eyes open), a subset of 1 to 12 maps (*k* = [1:12]) was randomly selected from the total set of voltage maps to serve as initial centroids for clustering. Spatial correlations between the *k* centroid maps and the remaining (unselected) voltage maps were then computed. These GFP maps were assigned to the centroid with which they had the highest spatial correlation, creating *k* clusters of maps. Correlation values were computed from the relative topographical configuration (but not the polarity) of the maps, and maps were only assigned to a cluster if the spatial correlation with the centroid map exceeded 0.5. After all correlations were calculated and individual maps assigned to a cluster, new centroid maps were created by averaging the constituent maps assigned to a given *k* cluster. Any remaining maps not assigned to a cluster were then compared to the recomputed (averaged) centroids and assigned again based on the correlation criterion. This process continued iteratively until the global explained variance (GEV) between the average centroids and the maps converged to a limit.

For each level of *k* = [1:12], this procedure was repeated 100 times, with a new subset of *k* centroids selected for each iteration. After 100 iterations, the set of centroids with maximal GEV were identified. For each individual recording, across all levels of *k*, the optimal number of *k* clusters was then selected from these sets of maximal GEV centroids, as determined using a metacriterion defined by 7 independent optimization criteria (see Custo et al., 2017; Bréchet et al., 2019). Figure 1 provides a schematic depicting the selection of voltage maps for clustering at GFP maxima from 1 second of eyes closed data from a random participant, and the result of the *k*-means clustering procedure for one full 8 minute recording.

**Figure 1:**
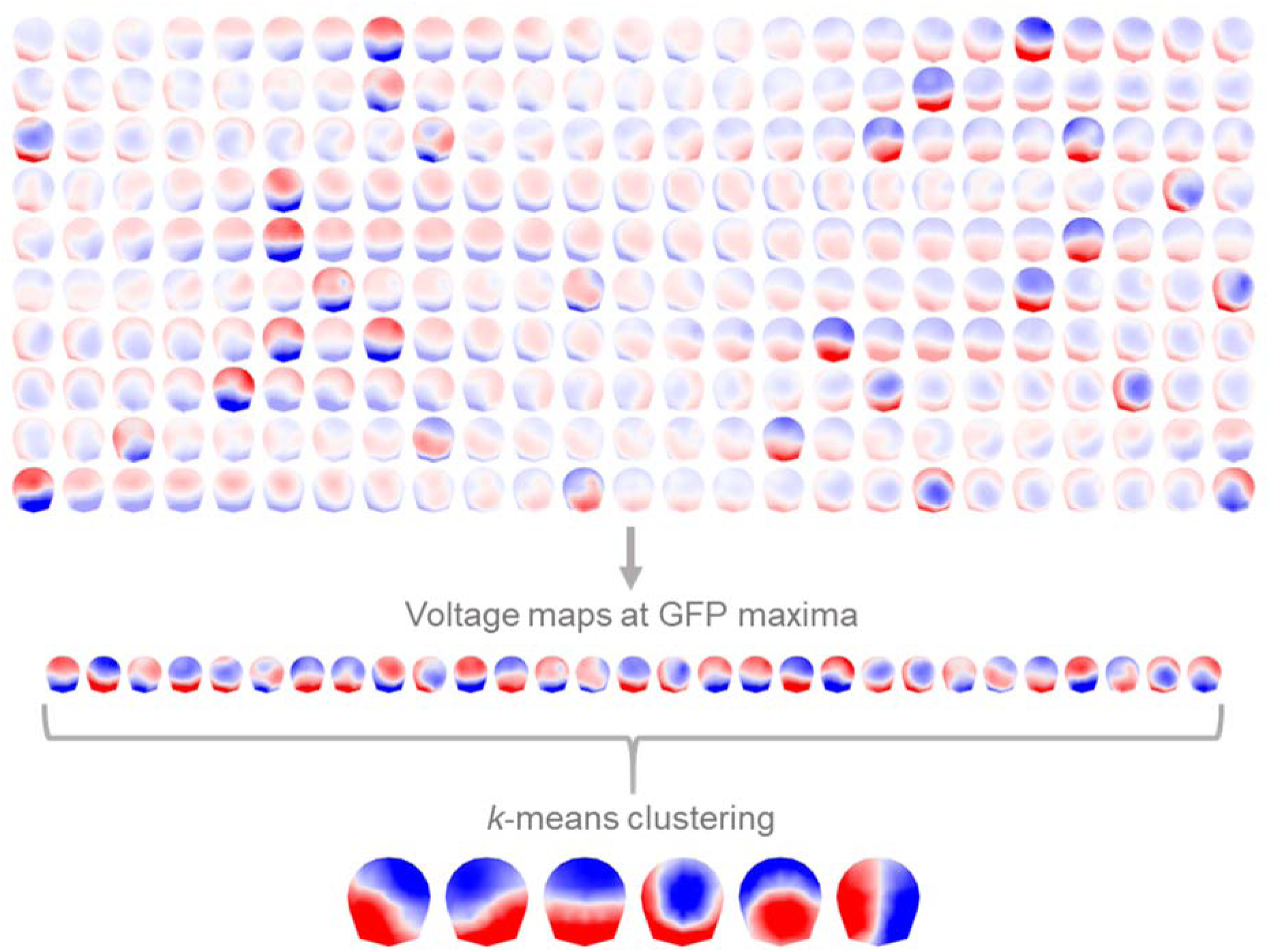
Rows depict the time series succession of voltage maps from left to right of 1 second of 64-channel eyes closed resting EEG (250 Hz sampling rate) from a recording chosen at random. Voltage maps are 2D isometric projections with nasion upwards. Voltage maps are identified at the local maxima in the global field power (GFP). The topography generally appears quasi-stable for several samples surrounding GFP peaks. *k*-means clustering of maps at GFP peaks (polarity is ignored) results in the optimal *k* clusters of voltage maps for the recording (8-minutes of EEG in total). The results of *k*-means clustering for this recording identified 6 cluster centroids that explained 85.93% of the topographic variance of maps at GFP maxima.

#### Clustering of subject-level centroids

In a second step, we conducted *k*-means clustering on the optimal centroids identified from the clustering of subject-level voltage maps just described. This was done to identify the optimal *global* clusters that best explained all subject-level representative cluster centroids. A set of *k* = [1:15] maps was randomly selected from the 382 subject-level topographies to use as centroids for clustering. For each level of *k*, 200 iterations were run, until the GEV converged to a limit and the *k* centroids with the maximal GEV were selected. Again, maps were only assigned if the spatial correlation exceeded 0.5.

After all iterations, the optimal number of *k* = [1:15] global clusters was determined using the optimization metacriterion, resulting in a set of *k* global centroids that best represent the overall subject-level topographic configurations.

#### Parameterization of the microstate time series

The global centroids were then fit back to the original EEG recordings to derive a time series sequences of microstates, separately for the eyes closed and open conditions. All samples of an individual subject’s continuous EEG were categorized by the global topography that demonstrated the highest spatial correlation between the sample-wise voltage map and the centroids of global clusters. EEG samples that had low spatial correlation (< .5) with all global centroids were left unassigned to a microstate centroid. Polarity was again ignored during centroid assignment. Temporal smoothing was applied to the continuous microstate sequence by ignoring segments that were present for less than six consecutive samples for a given centroid (< 24 msec), then assigning the first half of those brief segments to the preceding centroid and the second half to the subsequent centroids in the time series.

Three microstate parameters were derived from each subject-level microstate time series. *Global explained variance* (GEV), defined as the percentage of observed topographic variance explained by each specific global topographic centroid (i.e., microstate configuration). *Mean microstate duration*, or the average duration (in milliseconds) of continuous samples of the EEG time series categorized according to a specific microstate configuration. And *frequency of occurrence*, which represents how many times per second, on average, a given microstate occurred.

#### Transition probabilities and microstate sequence analysis

We also calculated first order Markov-chain transition probabilities from the time series of microstates using the *R* package *seqHMM* (Helske and Helske, 2019). The probability that a given microstate would transition to another microstate configuration was calculated for each pair of microstate configurations for each individual recording. Transitions to unassigned epochs lasting for more than 5 consecutive samples (> 20 msec) were excluded from calculations, while unassigned epochs shorter in duration were ignored and transitions were calculated based on the next occurring microstate.

### Individual Difference Assessment of Cognition, Personality, and Mood

#### Attentional performance

Three modules of the Test of Attentional Performance (TAP version 2.3.1; see Zimmermann and Fimm, 2002, and Babayan et al., 2019) were administered via computer to assess attentional performance. Errors, omissions, and reaction times (RTs) were recorded as outcomes of performance and provided as summary scores on the data-sharing repository. Two participants noted that they had forgotten their reading glasses during modules of the TAP. We nevertheless included these participants because their overall performance fell within acceptable range of their peers.

##### Alertness

In the psychomotor alertness TAP module, participants were asked to respond as quickly as possible to plus sign (“+”) stimuli presented on the monitor at randomly varying intervals. Stimuli were presented in four blocks under two conditions. In the first condition (blocks 1 and 4), participants simply responded as quickly as possible to presented stimuli. In the second condition (blocks 2 and 3), participants were also asked to respond as quickly as possible, but in this case the target stimuli were accompanied by an auditory warning signal that randomly sounded between 300 and 700 msec prior to when the target appeared. Mean reaction times (RT) for the no-signal and signal conditions were gathered for each participant. In addition, we computed the RT intra-individual coefficient of variability (ICV), calculated as the intra-individual RT standard deviation divided by the mean RT for each participant. Paired comparisons indicated that the mean RT in the no-signal condition (*M* = 231.98 msec, *SD* = 43.99) did not statistically differ (*p* = .755) from the mean RT in the signal condition (*M* = 232.55 msec, *SD* = 45.65). Similarly, ICV in the no-signal condition (*M* = 0.14, *SD* = 0.06) did not statistically differ (*p* = .462) from ICV in the signal condition (*M* = 0.15, *SD* = 0.06). The magnitude of the difference between conditions was negligible for RT (*d* = 0.023, 95% CI [− 0.226, 0.180]) and ICV (*d* = 0.054, 95% CI [−0.257, 0.149]). Accordingly, RTs and ICVs were each averaged across conditions to derive mean estimates reflecting participants’ task-averaged performance. Greater RT and ICV for an individual indicate slower and more variable speeded responding, respectively.

##### Response compatibility

In the response compatibility TAP module, participants were asked to respond as quickly as possible with the hand (left or right) that was congruent with the pointing direction, but not spatial location, of arrows presented either to the left or right of a central fixation point. That is, participants were asked to respond according to the *direction* of the arrow, irrespective of the side of the monitor on which the arrow appeared. Two participants were excluded because they responded incorrectly on nearly all trials (> 25 errors per condition). Reaction times were calculated separately for correct responses when the side of presentation was congruent with the direction of the arrow, and for correct responses when the side of presentation was incongruent with the pointing direction. Reaction times in the congruent condition (*M* = 445.54 msec, *SD* = 108.54) were significantly faster (*p* < .001, *d* = 0.579, 95% CI [0.371, 0.787]) than in the incongruent condition (*M* = 486.45, *SD* = 110.62). We quantified response compatibility as the difference in RT between the incongruent and congruent conditions for each participant. A larger response compatibility score indicates a greater response interference effect.

##### Working memory

In the working memory TAP module, participants viewed a series of numbers presented sequentially on the screen. They were asked to respond as quickly as possible whenever a number matched the number that was presented two trials previously. There were 15 series of numbers presented in total. One participant did not complete this module. Accuracy, in proportion correct, was calculated for each participant from the number of correctly identified stimuli.

#### Personality

Following the resting EEG recording, participants completed several self-report questionnaires assessing personality, motivation, affect, and emotionality (see Babayan et al., 2019). We selected two well-established measures of trait motivational tendencies and personality to explore as correlates of microstate parameters.

##### BIS-BAS

Participants completed the 24-item German translation (Strobel et al., 2001) of the Behavioral Inhibition and Behavioral Activation scale (BIS-BAS; Carver and White, 1994), to assess general tendencies towards avoiding aversive outcomes and approaching appetitive outcomes. The BIS-BAS assesses four motivational traits, including *punishment sensitivity* (BIS), relating to one’s reactions to the anticipation of punishment; *drive* (BAS Drive), or one’s tendency to persist in the pursuit of desired goals; *fun-seeking* (BAS Fun), or one’s tendency to desire new rewards and seek out fun situations; and *reward responsiveness* (BAS Reward), or one’s tendency to experience positive reactions to the occurrence or anticipation of rewards. Scale items were rated from 1 (“very false for me”) to 4 (“very true for me”), and summed to obtain subscale scores.

##### NEO-FFI

Participants completed the 60-item German translation (Borkenau and Ostendorf, 2008) of the NEO Five Factor Inventory (NEO-FFI; Costa and McCrae, 1992) to assess general personality traits. The NEO-FFI assessed 5 general traits encompassing *extraversion*, which includes the disposition to be talkative, energetic, and assertive; *neuroticism*, which includes being tense, moody, and anxious; *openness to experience*, including having broad interests and being imaginative and insightful; *agreeableness*, being sympathetic, kind, and affectionate; and *conscientiousness*, being organized, thorough, and reliable. Scale items were rated from 0 (“strongly disagree”) to 4 (“strongly agree”) and averaged for each personality dimension.

#### Mood

Immediately prior to the resting EEG recording, participants completed the 24-item Multidimensional Mood State Questionnaire (MDMQ; Steyer et al., 1997) to assess their current mood state. Mood was assessed from items associated with three dimensions: positive mood (*good to bad*); arousal (*awake to tired*); and anxiety (*calm to nervous*). Scale items were rated from 1 (“not at all”) to 5 (“very much”), and items were summed to obtain subscale scores. Lower scores represent participants feeling positive, awake, and calm. Data were missing from 3 individuals for measures of positive mood and anxiety, and from 5 individuals for arousal.

### Analysis

For within-subject analyses, we used linear mixed effects models to compare microstate parameters as function of microstate configuration (A, B, C, D, and E; see Results), perceptual condition (eyes closed and eyes open), and the interaction of these categorical fixed effects. Model parameters were estimated using restricted maximum likelihood in package *lme4* in *R* (Bates et al., 2015), and degrees of freedom were calculated by Satterthwaite approximation. Type III tests of fixed effects are reported for omnibus tests, and parameter estimates are given for models. The proportion of variance explained (*R*^2^) by the fixed effects was also calculated for each outcome (Nakagawa et al., 2017). Random subject intercepts were included to allow for within-person dependencies. In addition, for microstate duration and occurrence, we calculated the intraclass correlation (ICC) on a null (i.e., unconditional means) model using the *R* package *sjstats* (Lüdecke, 2019). For repeated measures data, the ICC can be used to describe the proportion of variance in each outcome that is attributable to between-subject versus within-subject differences (Hoffman, 2015). The ICC was not computed for the global explained variance because it is by definition scaled to account for differences in variability between individuals.

For between-subject analyses, independent samples comparisons were used to compare age groups (younger and older adults) and gender groups (female and male) on each microstate parameter. Standardized effect sizes were calculated as Cohen’s *d* with pooled variances using the *effsize* package in *R* (Torchiano, 2018); in select cases, *dz* was computed for paired comparisons (Gibbons et al., 1993). For correlational analyses, bivariate partial correlations were calculated between microstate parameters and measures of personality, mood, and attentional performance, controlling for age group differences. Resultant *p* values from these correlations were subjected to false discovery rate (FDR) control of Type I error (Benjamini and Hochberg, 1995). Microstate parameters were averaged across the eyes closed and eyes open conditions for all correlations and between-subject analyses.

Power analyses indicated that this study was strongly powered (1 – β = .95) with 191 participants to detect correlations of *r* > .25 in magnitude and standardized within-person differences of *d* > .26. Nevertheless, failing to identify statistically significant correlations provides no affirmative evidence in support of the null hypothesis that an effect is equal to zero. We therefore used a two one-sided test (TOST) equivalence testing procedure (Lakens, 2017) to evaluate evidence in favor of the null hypothesis. In the TOST procedure the null hypothesis is the presence of a minimally meaningful effect, and the alternative hypothesis is that the effect is weaker than the minimal effect size of interest. This is evaluated by comparing an observed effect with the lower (ΔL) and upper (ΔU) equivalence bounds of an effect deemed statistically meaningful using two one-sided comparisons. Here, a meaningful effect was defined as a correlation equal to or greater than *r* = ± .15 or a standardized mean difference of *d* = ± .3 or greater.

We also evaluated condition, age, and gender differences in transition probabilities for each microstate transition pair. Nonparametric permutation tests were used to account for Type I error (Nichols and Holmes, 2002) resulting from multiple comparisons of transition pairs. The strength of observed condition differences were each compared to a null distribution of the strongest effects occurring by chance among all the multiple comparisons of transition pairs for 10^4^ random permutations.

## Results

### Topographic Segmentation

The *k*-means clustering procedure revealed an optimal number of 4 to 8 subject-level centroid topographies for each individual EEG recording (*M* = 5.08, *SD* = 0.94, for a total of 1940 topographies across all recordings). In a second round of *k*-means clustering, five global clusters were identified that together explained 85.03% of the GEV in the 1940 individual subject-level cluster centroid topographies. These five clusters, which we designate as microstates A through E, were retained as the optimal number of global clusters based on the optimization metacriterion.

Figure 2 depicts the five optimal global cluster centroids and shows the 1938 individual subject-level cluster topographies grouped according to their global cluster membership. Two of the 1940 subject-level topographies went unassigned. It can be seen that these five global microstate configurations nicely represent the commonalities in topographic patterning among the individual participant recordings, irrespective of perceptual condition. To confirm this, we conducted a secondary *k*-means clustering of the 1940 subject-level centroid topographies, separately for the eyes closed and eyes open resting conditions. Figure 3 depicts the resultant global clusters and spatial topographic correlations from this analysis. Five optimal global clusters were identified for both the eyes closed and the eyes open conditions, which explained 85.48% and 85.12% of GEV in subject-level cluster centroids, respectively. Importantly, the five independently identified cluster topographies were essentially equivalent between conditions, with all topographic correlations exceeding, *r* = .99. Consequently, for all further analyses, we used the topographic configurations identified from the combined condition analyses, as illustrated in Figure 2.

**Figure 2:**
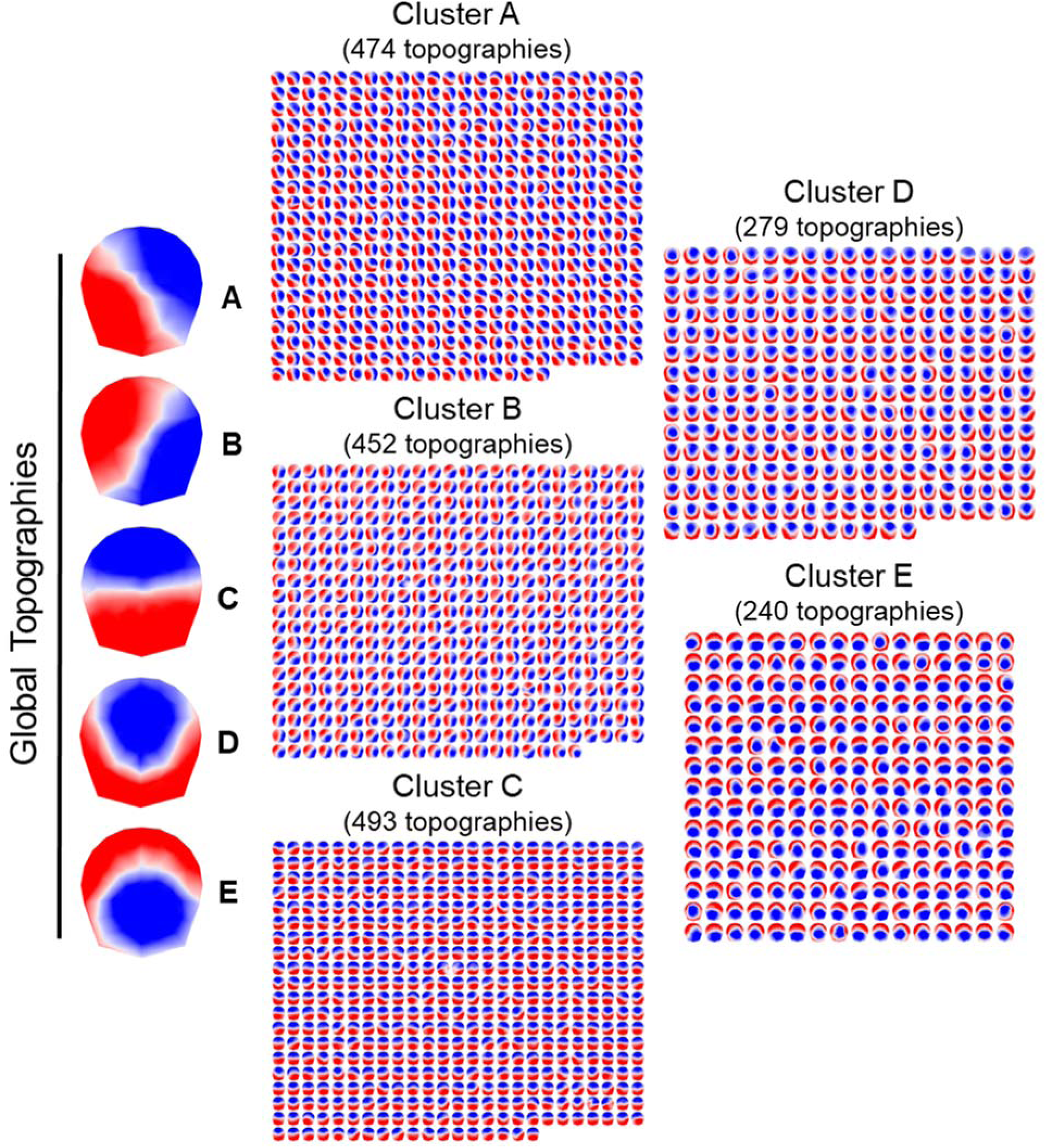
Five global cluster centroids were identified from *k*-means clustering during 8-min of eyes closed and 8-min of eyes open rest. 1938 cluster centroids derived from *k*-means clustering of 382 individual subject recordings are shown grouped according to their global cluster membership. Voltage topographies are 2D isometric projections with nasion upwards. Each global topography is the centroid of respective clusters of maps.

**Figure 3:**
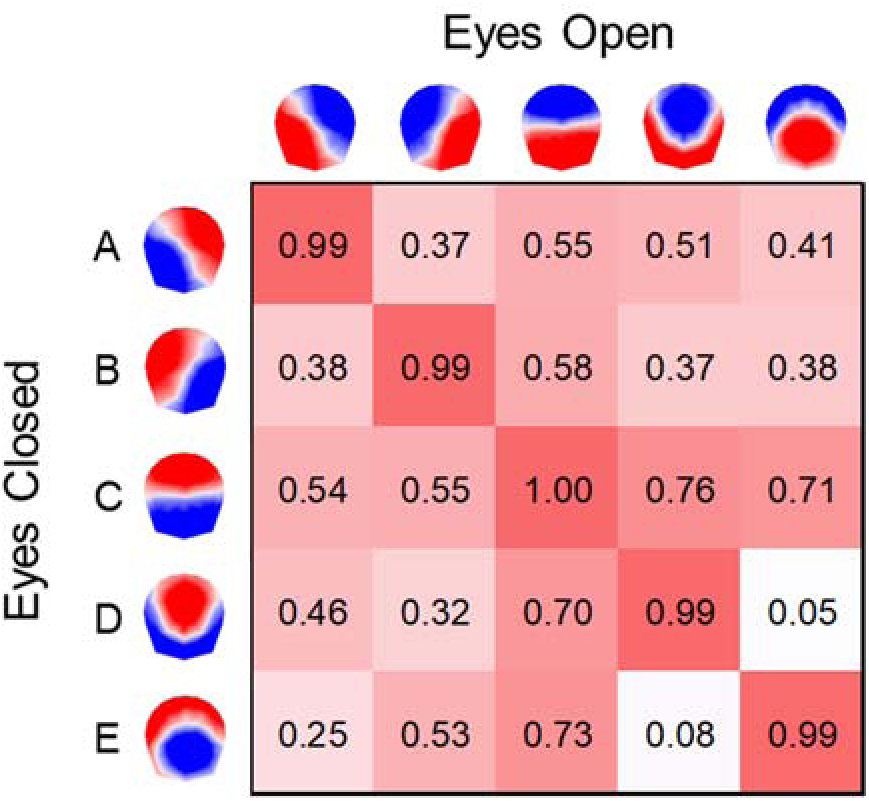
Spatial correlations are depicted between global microstate cluster topographies (A–E) identified from separate clustering of eyes closed and eyes open conditions. Polarity is ignored and only the spatial configuration is considered.

### Topographic Fitting and Microstate Analysis

The five global cluster centroids (A through E) were successfully assigned, on average, to 85.94% of samples taken from the continuous time series of the 382 original EEG recordings (*SD* = 11.11%). Four participants were excluded from all further analyses because a large percentage of voltage maps from their continuous EEG time series (> 45%) could not be assigned to a global cluster. Among the remaining 374 recordings, the five global microstate topographies together explained 65.03% (*SD* = 6.13%) of the GEV in the observed voltage maps in the eyes closed condition, and 60.99% (*SD* = 5.62%) of the GEV in the eyes open condition, on average across individuals. The effect of perceptual state on total GEV was significant (*b* = −4.044%, *p* < .001, 95% CI [−4.793, −3.295]), with differences between resting conditions explaining about 11% of the observed variance in total GEV (*R*^2^ = .106). Thus, the overall ability of microstates to explain the EEG time series was substantially influenced by an individual’s perceptual state at rest.

Once the global centroids were fit back to the observed topographies, the GEV, mean duration, and frequency of occurrence of each microstate configuration were calculated. Table 1 provides descriptive statistics for these parameters for the full EEG sample, and separately as a function of the eyes closed and eyes open conditions. The left hand side of Figure 4 plots the individual participant data along with the condition means and standard deviations. It can be seen from the figure that microstate C tended to predominate in the EEG time series for both perceptual conditions. Hence, for the within-subject models that follow, the categorical fixed effect of microstate configuration was centered at microstate C. The effect of perceptual condition was centered to the eyes closed state.

**Figure 4:**
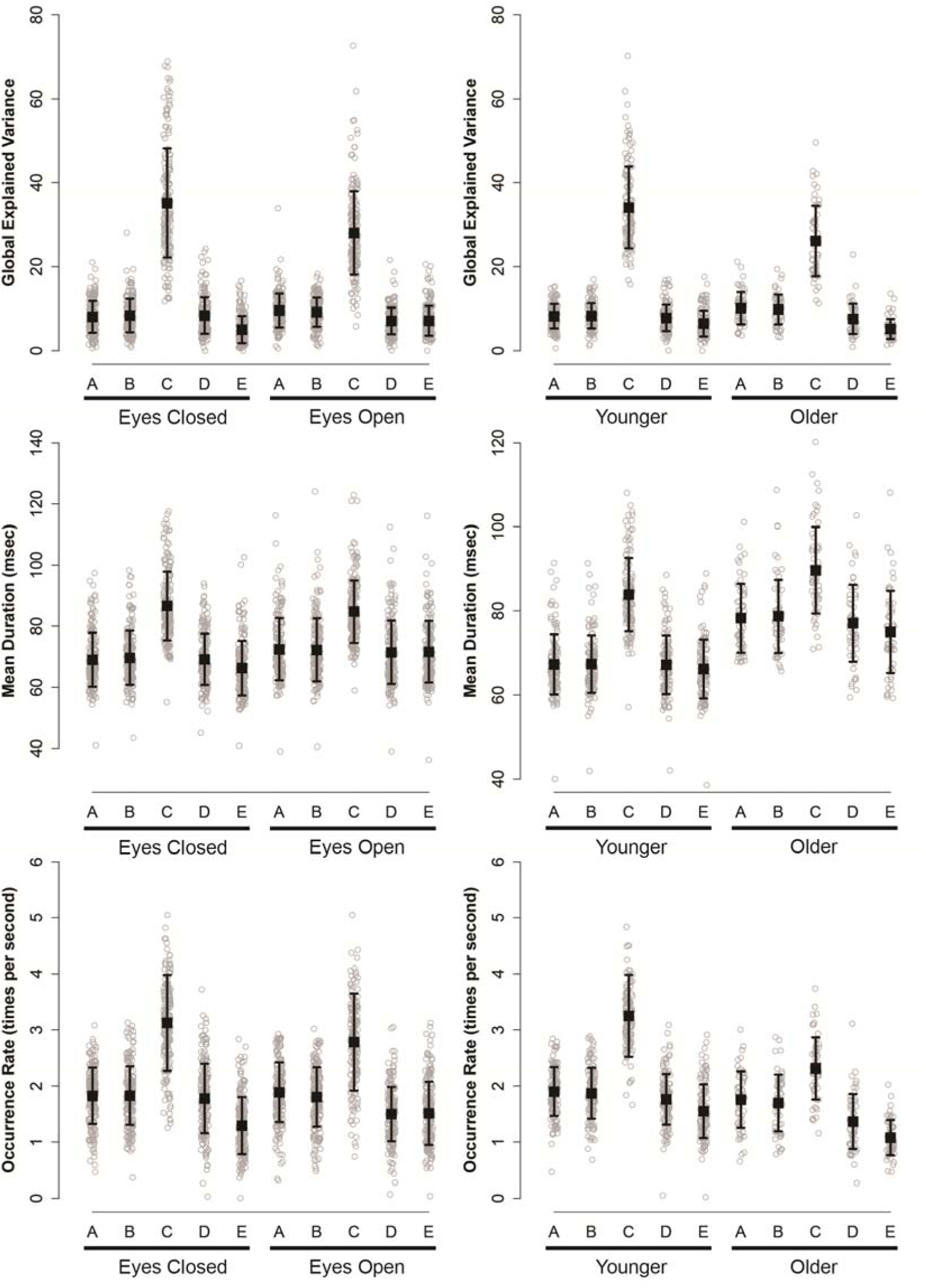
Means and standard deviations are plotted for global explained variance (GEV), microstate interval duration (in msec), and occurrence rate for each microstate configuration (A– E). Microstate parameters are provided for eyes closed and eyes open conditions, and age groups (younger *n* = 128, and older *n* = 59) collapsed across conditions. Observed subject averages (*n* = 187) are plotted as dots. Errors bars are the standard deviation around the mean.

**Table 1:**
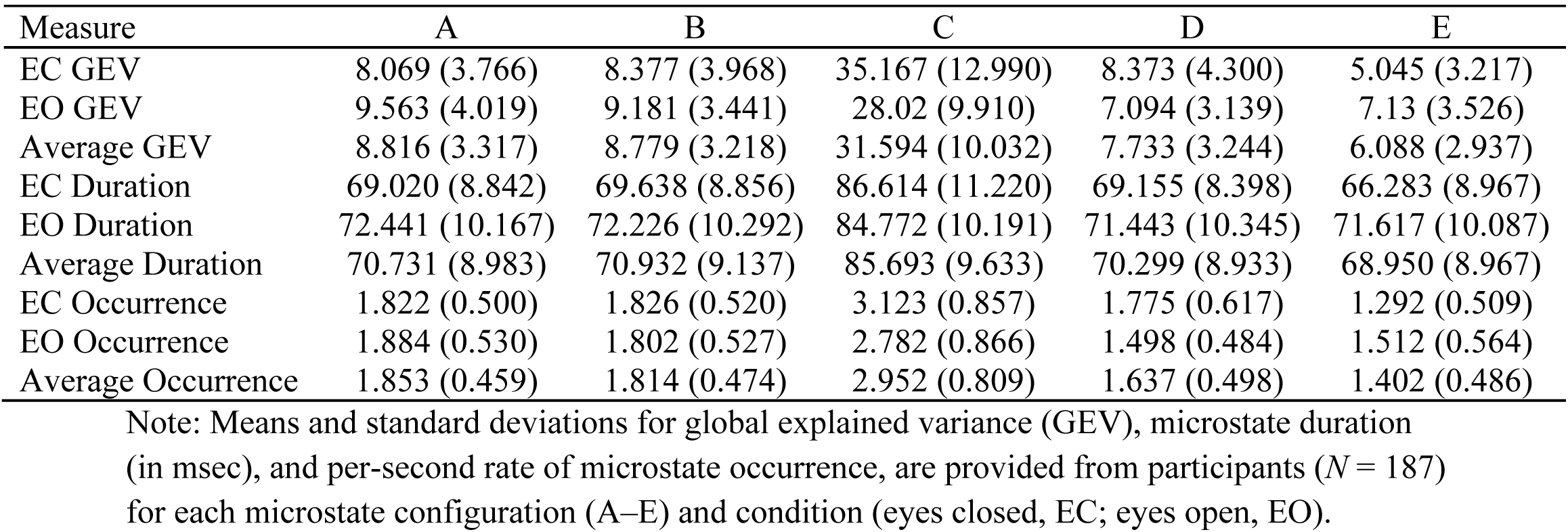
Descriptive statistics of microstate parameters from eyes closed (EC) and eyes open (EO) conditions.

#### Global explained variance

The mean percentage of observed topographic variance explained by each microstate configuration ranged from 5.05% to 35.17% in the eyes closed condition, and from 7.09% to 28.02% in the eyes open condition (see Table 1). However, microstate C was the only configuration that explained more than ten percent of the global topographic variance, on average, in either perceptual condition. In a mixed effects model, we observed a significant effect of microstate configuration, *F*(4, 1860) = 1133.36, *p* < .001, a significant effect of perceptual condition, *F*(1, 1860) = 8.13, *p* = .004, and a significant interaction between configuration and condition, *F*(4, 1860) = 35.23, *p* < .001. The parameter estimates from this model are reported in Table 2. For these parameters, the intercept reflects the mean GEV value for microstate C in the eyes closed condition. And, in order, the main effects of microstates A, B, D, and E, are the differences of these configurations from configuration C for the eyes closed condition; the eyes open effect is the difference between the eyes closed and open states for microstate C; and the interaction effects are the difference in how each microstate changes as a function of the eyes open condition relative to the change in microstate C.

**Table 2:** Model estimates from analyses of microstate parameters.

| Model Parameter | Estimate (SE) |  |  |
| --- | --- | --- | --- |
|  | GEV | Mean Duration | Occurrence |
| Fixed Effects |  |  |  |
| Intercept | 35.167 (0.448)*** | 86.614 (0.715)*** | 3.123 (0.045)*** |
| Microstate A | -27.098 (0.634)*** | -17.594 (0.566)*** | -1.301 (0.057)*** |
| Microstate B | -26.790 (0.634)*** | -16.976 (0.566)*** | -1.297 (0.057)*** |
| Microstate D | -26.794 (0.634)*** | -17.459 (0.566)*** | -1.348 (0.057)*** |
| Microstate E | -30.122 (0.634)*** | -20.331 (0.566)*** | -1.831 (0.057)*** |
| EO | -7.147 (0.634)*** | -1.841 (0.566)** | -0.342 (0.057)*** |
| EO x Microstate A | 8.641 (0.634)*** | 5.262 (0.800)*** | 0.404 (0.081)*** |
| EO x Microstate B | 7.951 (0.634)*** | 4.429 (0.800)*** | 0.317 (0.081)*** |
| EO x Microstate D | 5.868 (0.634)*** | 4.129 (0.800)*** | 0.065 (0.081) |
| EO x Microstate E | 9.232 (0.634)*** | 7.175 (0.800)*** | 0.562 (0.081)*** |
| Random Effects |  |  |  |
| Random Intercept | 0 | 65.6 (8.100) | 0.071 (0.2661) |
| Residual Variance | 37.6 (6.132) | 29.93 (5.471) | 0.305 (0.552) |
| Subjects (N) | 187 | 187 | 187 |
| Observations | 1870 | 1870 | 1870 |

Note: Maximum likelihood estimates are reported from models of global explained variance (GEV), microstate duration (in msec), and per-second rate of microstate occurrence, for fixed effects of microstate configuration (microstates A–E) and condition (EC and EO). Microstate C and the EC conditions serves as the reference. The number of subjects (*N*) and observations contributing to the analyses are provided. Standard errors are reported in parentheses. \* $p < .05$ , \*\* $p < .01$ , \*\*\* $p < .001$ .

As indicated in Table 2, microstates A, B, D, and E explained significantly less variance in the continuous EEG time series than configuration C (all *ps* < .001) when participants’ eyes were closed (see also Figure 4). When participants opened their eyes, however, the GEV for microstate C was reduced (*b* = −7.147%, *p* < .001, 95% CI [−8.387, −5.907]). Finally, the eyes open effect for microstates A, B, D, and E was found to differ in magnitude from that of microstate C (all *p*s < .001, for the interaction terms). For the simple effects of perceptual condition (not shown in in the table): Microstates A (*b* = 1.494%, *p* = .019, 95% CI [0.254, 2.734]) and E (*b* = 2.084%, *p* = .001, 95% CI [0.844, 3.325]) explained more variance in the eyes open than in the eyes closed condition, whereas microstate D explained less variance (*b* = - 1.279%, *p* = .044, 95% CI [−2.519, −0.039]). Microstate B, however, did not change between the eyes open and closed states (*b* = 0.084%, *p* = .205, 95% CI [−0.436, 2.044]).

#### Mean duration

On average, each microstate occurrence lasted between 66.28 and 86.61 msec in the eyes closed state, and between 71.44 and 84.77 msec in the eyes open state (from Table 1). The intra-class correlation for mean duration was .446, suggesting that 44.6% of variance in mean microstate duration was attributable to differences between individuals, but that the majority of variance (55.4%), was attributable to within-person variation. Comparison across microstates and conditions revealed a significant effect of microstate configuration, *F*(4, 1674) = 605.18, *p* < .001, a significant effect of perceptual condition, *F*(1, 1674) = 86.84, *p* = .004, and a significant interaction between configuration and condition, *F*(4, 1674) = 21.62, *p* < .001. When considered alone, microstate configuration explained 28% of the observed outcome variance (*R*^2^ = .283) in mean duration, while all the significant fixed effects together explained 30% of the overall variance (*R*^2^ = .303).

Table 2 presents the parameter estimates for within-subject differences in mean microstate duration. Microstates A, B, D, and E were significantly shorter, on average, than configuration C (all *ps* < .001) in the eyes closed condition. However, the duration of configuration C shortened when participants opened their eyes (*b* = −1.841 msec, *p* = .001, 95% CI [−2.948, −0.735]), whereas the duration of microstates A (*b* = 3.421 msec, *p* < .001, 95% CI [2.315, 4.528]), B (*b* = 2.589 msec, *p* < .001, 95% CI [1.481, 3.694]), D (*b* = 2.288 msec, *p* < .001, 95% CI [1.181, 3.395]), and E (*b* = 5.334 msec, *p* < .001, 95% CI [4.227, 6.440]) were all significantly longer in the eyes open condition.

#### Occurrence

Microstates A, B, D, and E all occurred between one and two times per second per condition, on average, whereas configuration C occurred about 3 times per second overall (Table 1). The ICC for microstate occurrence was .056, such that only 5.6% of the variance in this measure could be attributed to differences between individuals. The preponderance of variance (94.4%) was instead attributable to within-person variation. As with GEV and microstate duration, we observed a significant effect of microstate configuration, *F*(4, 1674) = 438.55, *p* < .001, a significant effect of perceptual condition, *F*(1, 1674) = 7.95, *p* = .005, and a significant interaction between these effects, *F*(4, 1674) = 16.91, *p* < .001. The effect of microstate configuration alone explained 43% of the outcome variance (*R*^2^ = .425) in mean occurrence rate; together, all fixed effects explained 44% of the overall variance (*R*^2^ = .443).

The parameter estimates of Table 2 show that microstates A, B, D, and E again all differed from configuration C in the eyes closed condition (all *ps* < .001), and that microstate C occurred less frequently, on average, when participants had their eyes open (*b* = −0.342 times per second, *p* < .001, 95% CI [−0.453, −0.230]). Finally, the magnitude of change between perceptual conditions for microstates A, B, and E differed from the observed change for microstate C (all *p*s < .001, for the interaction terms). Simple effect comparisons showed that microstates A (*b* = 0.062, *p* = .276, 95% CI [−0.049, 0.174]) and B (*b* = −0.024, *p* = .671, 95% CI [−0.136, 0.087]) did not differ between conditions, whereas microstate D was significantly less frequent (*b* = - 0.277, *p* < .001, 95% CI [−0.388, −0.165]) and microstate E more frequent (*b* = 0.220, *p* < .001, 95% CI [0.109, 0.332]) in the eyes open condition.

### Individual Difference Correlates of Microstate Parameters

We next conducted between-subject analyses on microstate GEV, duration, and occurrence. Table 3 presents means and standard deviations for the predictors of personality, mood, and attentional performance for the final EEG sample of 187 participants (see Topographic Fitting and Microstate Analysis). Effect size differences between the younger and older age groups are also provided for each individual difference measure. Older adults were less neurotic, less open to experience, more conscientious, less fun seeking, and less inhibited, overall. They also reported a more fatigued or tired mood state than did younger adults. The largest effect sizes were observed for the attentional performance measures, in which older adults had slower RTs, were more variable in their responding, had a larger response compatibility effect, and demonstrated poorer working memory accuracy.

**Table 3:** Descriptive statistics for age, personality, mood, and attentional performance.

| Variable | Total sample |  |  | Younger adults |  | Older adults |  | Cohen's d |
| --- | --- | --- | --- | --- | --- | --- | --- | --- |
|  | *M* | *SD* | *N* | *M* | *SD* | *M* | *SD* |  |
| Personality |  |  |  |  |  |  |  |  |
| BIS | 19.89 | 3.13 | 187 | 20.34 | 3.23 | 18.92 | 2.67 | -0.466** |
| BAS Drive | 11.95 | 2.11 | 187 | 11.85 | 2.16 | 12.17 | 1.98 | 0.151 |
| BAS Reward | 16.95 | 1.97 | 187 | 17.06 | 1.91 | 16.69 | 2.08 | -0.187 |
| BAS Fun | 12.22 | 1.69 | 187 | 12.49 | 1.64 | 11.64 | 1.66 | -0.516** |
| Neuroticism | 1.48 | 0.55 | 187 | 1.55 | 0.58 | 1.33 | 0.46 | -0.407* |
| Extraversion | 2.43 | 0.51 | 187 | 2.44 | 0.54 | 2.40 | 0.43 | -0.090 |
| Openness | 2.70 | 0.50 | 187 | 2.78 | 0.52 | 2.53 | 0.41 | -0.525** |
| Agreeableness | 2.76 | 0.44 | 187 | 2.76 | 0.44 | 2.78 | 0.44 | 0.037 |
| Conscientiousness | 2.67 | 0.60 | 187 | 2.52 | 0.61 | 3.02 | 0.43 | 0.891*** |
| Mood |  |  |  |  |  |  |  |  |
| Good–Bad | 34.63 | 3.97 | 184 | 34.40 | 3.57 | 35.12 | 4.73 | 0.182 |
| Awake–Tired | 31.03 | 6.38 | 182 | 30.00 | 6.43 | 33.36 | 5.67 | 0.541*** |
| Calm–Nervous | 33.41 | 4.57 | 184 | 33.45 | 4.30 | 33.33 | 5.16 | -0.027 |
| Attention performance |  |  |  |  |  |  |  |  |
| Alertness RT | 232.27 | 43.1 | 187 | 217.25 | 29.4 | 264.86 | 49.8 | 1.286*** |
| Alertness ICV | 0.14 | 0.05 | 187 | 0.13 | 0.04 | 0.17 | 0.05 | 0.707*** |
| Compatibility | 42.50 | 72.4 | 185 | 27.54 | 49.5 | 75.28 | 99.6 | 0.691*** |
| Working memory | 0.87 | 0.15 | 186 | 0.91 | 0.11 | 0.79 | 0.20 | -0.873*** |

Note: Means and standard deviations are provided for age in years, measures of personality, mood, and attentional performance. *N* denotes complete sample size for each measure.
Descriptive statistics are given for both the younger ( $n = 128$ ) and older ( $n = 59$ ) age groups, and standardized mean differences (Cohens *d*) for comparisons between younger and older age groups are indicated. \* $p < .05$ , \*\* $p < .01$ , \*\*\* $p < .001$ .

#### Age and gender

We first examined differences between age groups and between genders through independent samples comparisons. Table 4 summarizes the effect sizes for these comparisons for each microstate configuration (averaged across the eyes open and eyes open conditions).

**Table 4:** Standardized effect sizes between age and gender groups.

| Measure | A | B | C | D | E |
| --- | --- | --- | --- | --- | --- |
| GEV |  |  |  |  |  |
| Age | .596*** | .479** | -.854*** | -.068 | -.451** |
| Gender | .026† | .040† | .199 | .112 | -.046† |
| Mean Duration |  |  |  |  |  |
| Age | 1.484*** | 1.517*** | .622*** | 1.275*** | 1.103*** |
| Gender | -.345* | -.400** | -.258 | -.349* | -.421** |
| Occurrence |  |  |  |  |  |
| Age | -.316* | -.364** | -1.372*** | -.855*** | -1.090*** |
| Gender | .333* | .409** | .321* | .291 | .184 |

Note: Standardized mean differences ( $d$ ) for comparisons between age and gender groups are reported for microstate parameters separately for each microstate configuration (A–E) averaged across conditions. Positive standardized mean differences reflect higher values for older and male participants, respectively. Significance values are indicated from independent samples comparisons. † = equivalent to zero, \* $p < .05$ , \*\* $p < .01$ , \*\*\* $p < .001$ .

All but one of the comparisons between age groups were significant. Older individuals had greater GEV for microstates A (*d* = 0.596, *p* < .001, 95% CI [0.280, 0.912]) and B (*d* = 0.479, *p* = .003, 95% CI [0.165, 0.793]), and less GEV for microstates C (*d* = −0.854, *p* < .001, 95% CI [−1.176, −0.531]) and E (*d* = −0.451, *p* = .005, 95% CI [−0.764, −0.137]). Mean duration was longer for older individuals for all microstate configurations: A (*d* = 1.484, *p* < .001, 95% CI [1.139, 1.830]), B (*d* = 1.517, *p* < .001, 95% CI [1.170, 1.864]), C (*d* = 0.622, *p* < .001, 95% CI [0.305, 0.939]), D (*d* = 1.275, *p* < .001, 95% CI [0.938, 1.611]), and E (*d* = 1.103, *p* < .001, 95% CI [0.773, 1.433]). Finally, older individuals had fewer occurrences of microstates, on average, for all microstate configurations: A (*d* = −0.316, *p* = .046, 95% CI [−0.628, −0.003]), B (*d* = - 0.364, *p* = .022, 95% CI [−0.677, −0.051]), C (*d* = −1.372, *p* < .001, 95% CI [−1.713, −1.032]), D (*d* = −0.855, *p* < .001, 95% CI [−1.177, −0.532]), and E (*d* = −1.090, *p* < .001, 95% CI [−1.420, −0.760]). Figure 4 (right hand side) plots the individual participant data for these comparisons, along with the means and standard deviations for each age group.

No significant differences were found for GEV as a function of gender. Moreover, for microstates A, B, and E, the effect of gender was found to be statistically equivalent to zero. Males had shorter mean durations of microstates A (*d* = −0.345, *p* = .025, 95% CI [−0.648, - 0.042]), B (*d* = −0.400, *p* = .009, 95% CI [−0.703, −0.096]), D (*d* = −0.349, *p* = .023, 95% CI [−0.652, −0.046]), and E (*d* = −0.421, *p* = .006, 95% CI [−0.725, −0.118]). Males also had more frequent occurrences of microstates A (*d* = 0.333, *p* = .030, 95% CI [0.030, 0.635]), B (*d* = 0.409, *p* = .008, 95% CI [0.105, 0.713]), and C (*d* = 0.321, *p* = .037, 95% CI [0.018, 0.623]).

#### Personality, mood, and attentional performance

We next examined bivariate correlations between each individual difference predictor and each microstate parameter. Because age group differences were observed for many of the predictor variables (see Table 3), we controlled for age group in all correlations and report partial correlation coefficients (as *r* statistics) for the overall sample.

##### Global explained variance

Table 5 presents the partial correlations for microstate GEV. Several correlations were identified as significant (*p*s < .05), however none of these remained significant following FDR control for 80 statistical tests. We report these correlations below but caution that a number of these effects may reflect Type 1 error. Furthermore, 25 correlations were identified as statistically equivalent to zero. These were correlations that ranged between *r* = -.03 and *r* = .03, and were significantly smaller than the lower (ΔL *r* = −.15) or upper (ΔU *r* = .15) equivalence bounds. The remaining correlations were statistically undetermined.

**Table 5:** Correlations between microstate GEV and measures of personality, mood, and cognition.

| Measure | GEV |  |  |  |  |
| --- | --- | --- | --- | --- | --- |
|  | A | B | C | D | E |
| Personality |  |  |  |  |  |
| BAS Drive | -0.104 | -0.152* | 0.018† | 0.089 | 0.105 |
| BAS Reward | -0.076 | -0.086 | -0.009† | 0.051 | 0.077 |
| BAS Fun | 0.003† | 0.134 | 0.015† | -0.123 | -0.077 |
| BIS | -0.022† | 0.043 | 0.072 | -0.038 | -0.098 |
| Neuroticism | -0.061 | 0.010† | 0.129 | -0.144 | -0.099 |
| Extraversion | 0.005† | -0.034 | -0.104 | 0.174* | 0.067 |
| Openness | 0.043 | -0.003† | -0.093 | 0.138 | -0.026† |
| Agreeableness | -0.010† | 0.078 | -0.068 | 0.177* | 0.006† |
| Conscientiousness | -0.115 | -0.215** | -0.003† | 0.186* | 0.181* |
| Mood |  |  |  |  |  |
| Good - Bad | -0.015† | -0.012† | 0.041 | 0.018† | -0.114 |
| Awake - Tired | -0.055 | 0.025† | 0.098 | -0.063 | -0.096 |
| Calm - Nervous | -0.018† | 0.002† | 0.047 | 0.040 | -0.168* |
| Attention Performance |  |  |  |  |  |
| Alertness RT | -0.036 | -0.097 | 0.009† | 0.121 | 0.099 |
| Alertness ICV | -0.137 | -0.095 | 0.041 | 0.079 | 0.106 |
| Compatibility | 0.005† | -0.045 | 0.010† | -0.139 | 0.063 |
| Working Memory | -0.045 | 0.069 | -0.001† | 0.080 | -0.026† |

Note: Partial correlation coefficients controlling for age group are reported for measures of personality, mood, and attentional performance for microstate global explained variance (GEV) averaged across conditions separately for each microstate configuration (A–E). † = equivalent to zero, \* $p < .05$ , \*\* $p < .01$ , \*\*\* $p < .001$ .

Drive was negatively correlated with GEV for microstate B (*r* = −0.152, *p* = .037, 95% CI [−0.290, −0.009]). Extraversion was positively correlated with microstate D (*r* = 0.174, *p* = .017, 95% CI [0.031, 0.310]), as was agreeableness (*r* = 0.177, *p* = .015, 95% CI [0.034, 0.313]). Conscientiousness was negatively correlated with microstate B (*r* = −0.215, *p* = .003, 95% CI [− 0.348, −0.074]), and positively correlated with microstates D (*r* = 0.186, *p* = .011, 95% CI [0.044, 0.321]), and E (*r* = 0.181, *p* = .014, 95% CI [0.038, 0.319]). Finally, nervous mood was negatively correlated with GEV for microstate E (*r* = −0.168, *p* = .023, 95% CI [−0.306, −0.025]).

##### Mean duration

Table 6 summarizes the partial correlations for microstate duration. Twelve significant correlations were identified (*p*s < .05), but none survived FDR control. There were also 16 correlations found equivalent to zero.

**Table 6:** Correlations between microstate duration and measures of personality, mood, and cognition.

| Measure | Duration |  |  |  |  |
| --- | --- | --- | --- | --- | --- |
|  | A | B | C | D | E |
| Personality |  |  |  |  |  |
| BAS Drive | -0.009† | -0.063 | -0.026† | 0.049 | 0.049 |
| BAS Reward | 0.072 | 0.031 | 0.022† | 0.102 | 0.095 |
| BAS Fun | 0.059 | 0.083 | 0.036 | -0.018† | 0.020† |
| BIS | -0.031 | -0.001† | 0.029† | -0.076 | -0.091 |
| Neuroticism | 0.102 | 0.143 | 0.178* | 0.023† | 0.051 |
| Extraversion | -0.001† | -0.029† | -0.119 | 0.054 | 0.020† |
| Openness | 0.016† | -0.011† | -0.063 | 0.071 | 0.027† |
| Agreeableness | -0.188* | -0.186* | -0.200** | -0.110 | -0.163* |
| Conscientiousness | -0.130 | -0.194** | -0.128 | 0.017† | -0.011† |
| Mood |  |  |  |  |  |
| Good - Bad | -0.007† | -0.026† | 0.013† | -0.016† | -0.047 |
| Awake - Tired | -0.005† | -0.014† | 0.064 | -0.060 | -0.042 |
| Calm - Nervous | 0.014† | 0.003† | 0.036 | -0.001† | -0.051 |
| Attention Performance |  |  |  |  |  |
| Alertness RT | -0.170* | -0.191** | -0.127 | -0.100 | -0.111 |
| Alertness ICV | -0.052 | -0.043 | -0.011† | 0.029† | 0.028† |
| Compatibility | 0.192** | 0.149* | 0.181* | 0.125 | 0.197** |
| Working Memory | -0.114 | -0.077 | -0.131 | -0.077 | -0.119 |

Note: Partial correlation coefficients controlling for age group are reported for measures of personality, mood, and attentional performance for mean microstate duration averaged across conditions separately for each microstate configuration (A–E). † = equivalent to zero, \* $p < .05$ , \*\* $p < .01$ , \*\*\* $p < .001$ .

For the personality measures, neuroticism was positively correlated with duration for microstate C (*r* = 0.178, *p* = .015, 95% CI [0.035, 0.313]), and agreeableness was negatively correlated microstates A (*r* = −0.188, *p* = .010, 95% CI [−0.323, −0.046]), B (*r* = −.186, *p* = 0.011, 95% CI [−0.321, −0.044]), C (*r* = −0.200, *p* = .006, 95% CI [−0.334, −0.059]), and E (*r* = −0.163, *p* = .026, 95% CI [−0.299, −0.020]). In addition, conscientiousness was negatively correlated with duration for microstate B (*r* = −0.194, *p* = .008, 95% CI [−0.329, −0.052]). Correlations were also observed for the attentional performance measures. Alertness RT was negatively correlated with microstates A (*r* = −0.170, *p* = .020, 95% CI [−0.306, −0.027]) and B (*r* = −0.191, *p* = .009, 95% CI [−0.326, −0.049]), and response compatibility was positively correlated with microstates A (*r* = 0.192, *p* = .009, 95% CI [0.051, 0.329]), B (*r* = 0.149, *p* = .044, 95% CI [0.007, 0.289]), C (*r* = 0.181, *p* = 0.014, 95% CI [0.040, 0.319]), and E (*r* = 0.197, *p* = .007, 95% CI [0.057, 0.335]).

##### Occurrence

Table 7 provides the partial correlations for microstate occurrence. There were 11 significant correlations (*p*s < .05), but once again, none survived FDR control. Twenty-one correlations were found to be equivalent to zero.

**Table 7:** Correlations between microstate occurrence and measures of personality, mood, and cognition.

| Measure | Occurrence |  |  |  |  |
| --- | --- | --- | --- | --- | --- |
|  | A | B | C | D | E |
| Personality |  |  |  |  |  |
| BAS Drive | -0.112 | -0.158* | 0.041 | 0.061 | 0.081 |
| BAS Reward | -0.081 | -0.100 | -0.015† | 0.036 | 0.047 |
| BAS Fun | -0.021† | 0.091 | -0.019† | -0.117 | -0.058 |
| BIS | -0.014† | 0.049 | 0.059 | -0.026† | -0.076 |
| Neuroticism | -0.106 | -0.062 | 0.047 | -0.157* | -0.140 |
| Extraversion | 0.041 | -0.018† | -0.054 | 0.183* | 0.081 |
| Openness | 0.019† | -0.031 | -0.032 | 0.089 | -0.002† |
| Agreeableness | 0.081 | 0.167* | 0.067 | 0.216** | 0.086 |
| Conscientiousness | -0.077 | -0.146 | 0.096 | 0.200** | 0.201** |
| Mood |  |  |  |  |  |
| Good - Bad | -0.006† | -0.005† | -0.018† | -0.008† | -0.092 |
| Awake - Tired | -0.029 | 0.043 | -0.031 | -0.045 | -0.066 |
| Calm - Nervous | -0.036 | -0.009† | -0.014† | 0.006† | -0.148* |
| Attention Performance |  |  |  |  |  |
| Alertness RT | 0.022† | 0.009† | 0.083 | 0.194** | 0.159* |
| Alertness ICV | -0.103 | -0.074 | 0.019† | 0.100 | 0.101 |
| Compatibility | -0.095 | -0.131 | -0.090 | -0.153* | 0.003† |
| Working Memory | 0.021† | 0.075 | 0.067 | 0.080 | -0.014† |

Note: Partial correlation coefficients controlling for age group are reported for measures of personality, mood, and attentional performance for mean microstate occurrence rate separately for each microstate configuration (A–E) averaged across conditions. † = equivalent to zero, \* $p < .05$ , \*\* $p < .01$ , \*\*\* $p < .001$ .

Drive was negatively correlated with occurrence for microstate B (*r* = −0.158, *p* = .031, 95% CI [−0.295, −0.015]). Neuroticism was negatively correlated with microstate D (*r* = −0.157, *p* = .033, 95% CI [−0.294, −0.013]), whereas extraversion was positively correlated with microstate D (*r* = 0.183, *p* = .012, 95% CI [0.040, 0.318]). In addition, agreeableness was positively correlated with microstates B (*r* = 0.167, *p* = .023, 95% CI [0.024, 0.303]) and D (*r* = 0.216, *p* = .003, 95% CI [0.075, 0.348]), and conscientiousness was positively correlated with microstates D (*r* = 0.200, *p* = .006, 95% CI [0.058, 0.334]) and E (*r* = 0.201, *p* = .006, 95% CI [0.059, 0.335]). A negative correlation was also observed between anxious mood and microstate E (*r* = −0.148, *p* = .045, 95% CI [−0.288, −0.005]). Finally, for attentional performance, alertness RT was positively correlated with occurrence for microstates D (*r* = .194, *p* = .008, 95% CI [0.052, 0.328]) and E (*r* = .159, *p* = .030, 95% CI [0.016, 0.296]), and response compatibility was negatively correlated with occurrence for microstate D (*r* = −.153, *p* = .038, 95% CI [−0.291, - 0.009]).

### Perceptual State Differences in Microstate Transition Dynamics

In a final analysis, we examined whether the probability of transitioning from one microstate configuration to another differed between the eyes closed and eyes open conditions. The set of five topographic configurations (A–E) allowed for 20 pairs of Markov-chain transition probabilities. Figure 5 presents these transition probabilities for each condition, as well as the standardized mean differences between conditions for each transition pair. Significant differences between the eyes closed and open conditions were found for 14 pairs of microstate transitions (all *p*s < .041). The standardized mean differences for these significant comparisons ranged from small to moderate (*dz* range = .151 to .631) in magnitude. However, of these, only differences larger than *dz* = .31 could be considered significantly stronger than expected under the null hypothesis after permutation-based correction for multiple comparisons.

**Figure 5:**
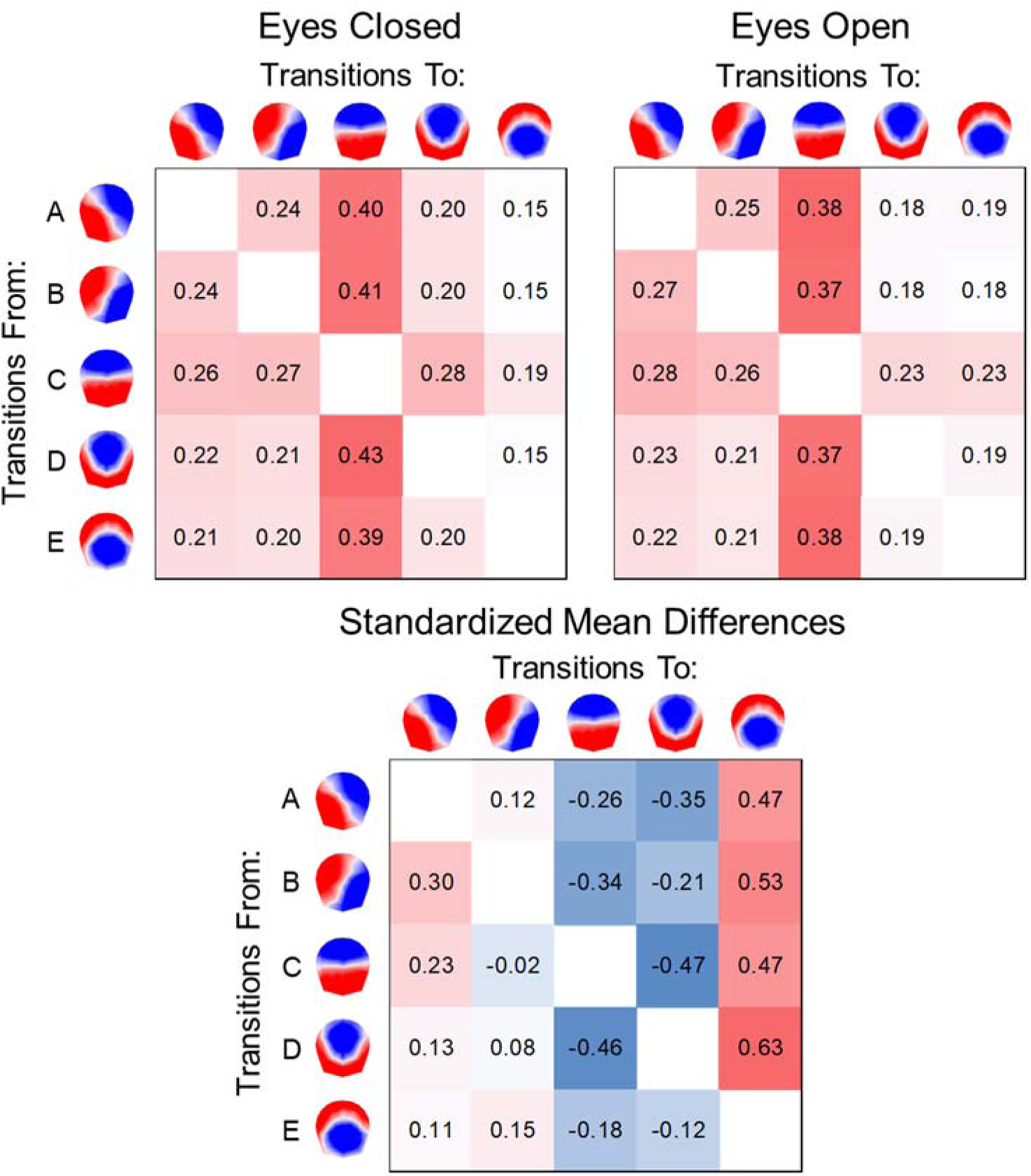
Mean Markov-chain transition probabilities for resting state microstates microstate configuration (A–E) are shown (top) based on 187 sets of microstate sequences for the eyes closed and eyes open conditions. The probabilities of transitions are depicted from each microstate on the vertical axis to microstates on the horizontal axis. Standardized mean differences (*dz*) for comparisons between conditions are indicated (bottom) for all transition pairs.

No significant differences in transition dynamics were observed between age groups (*d* range = −.188 to .259, *p*s > .101), or between genders (*d* range = −.275 to .282, *p*s > .066) for any transition pairs (when averaged across perceptual conditions).

## Discussion

The findings of this study show that the dynamics of ongoing brain electrocortical activity are sensitive to within-person differences in patterns of scalp-recorded voltage topography. Neuroelectric activity recorded during eyes-closed rest and during simple visual fixation was segmented into a time series of transient microstate intervals. In general, resting perceptual state (eyes closed vs. open) was seen to be the strongest determinant of microstate temporal dynamics, and was the only factor that systematically influenced transition probabilities between successive microstate configurations in individuals over time. However, our findings also offer robust exploratory evidence, from a well-powered sample, that microstate parameters can differentiate groups based on gender and age and may be weakly to moderately associated with individual differences in personality and cognition.

It was found that five data-driven microstate configurations (designated A through E) could account for the preponderance of variance in the EEG time series of the 374 recordings (from 187 participants) included in the study. Moreover, the temporal dynamics associated with these microstates were found to predominately vary within individuals. This was demonstrated by a simple portioning of the unexplained variance in mean microstate duration and occurrence rate into its within-person and between-person components; and was further confirmed by the fact that about 30 to 44% of the total outcome variance in these parameters could be explained by the within-person factors of perceptual condition and microstate configuration. On the basis of these estimates, we can conclude that individuals varied within themselves to an equal or greater degree than they differed between each other, and recommend that researchers turn more of their attention towards understanding the factors that contribute to within-person variation in the activity of whole-brain neuronal networks.

Resting perceptual state was an important source of change in microstate temporal dynamics. The most consistent of these effects was found for configurations C and E. When participants opened their eyes, microstate C explained less variance in the ongoing EEG time series, was of shorter duration, and occurred less frequently than when their eyes were closed. Microstate E, by contrast, exhibited the opposite pattern, increasing in variance explained and in duration and occurrence in the eyes open state. Notably, these effects were also mirrored by changes in transition probabilities. The transitions of B → C, C → D, and D → C (as well as A → D) were less likely to occur, while transitions to microstate E were more likely to occur when individuals opened their eyes. The patterns for configurations A, B, and D were less uniform across outcomes; though all configurations other than C increased in duration when participants opened their eyes. Overall, the differential effects of perceptual state on microstate configurations—some increasing and others decreasing their predominance in the time series— could reflect the operation of compensatory network dynamics, reflecting variation in cognitive operations between states. When eyes are open, for instance, the shortened duration and reduced occurrence of microstate C appears to be offset by increases in the duration and prevalence of other microstates. Compensatory shifts of this kind, from internally focused (eyes closed) to more externally focused (eyes open) perceptual states, are reminiscent of the functional competition observed between the default mode and attention and sensory networks with fMRI (Raichle, 2015).

Our findings partially conflict with those of Seitzman et al. (2017), who compared microstate parameters between brief (25 second) periods of eyes closed and eyes open rest in a sample of 24 participants. They reported significant reductions in microstate D duration and increases in occurrence rate for microstate B when participants opened their eyes. They also found no support for differences in microstate transition dynamics between perceptual states. In contrast, we observed increases in microstate D duration and no change in the occurrence rate of microstate B when participants opened their eyes, as well as systematic change in microstate transition dynamics between conditions. Reductions in total GEV in the eyes open condition, however, was consistent between studies. The discrepancies between their study and our own could have arisen due to differences in how the resting state was operationalized. Seitzman and colleagues (2017) instructed their participants to let their minds freely wander when at rest, whereas Babayan et al. (2019), whose data we examined, simply asked participants to rest quietly during the eyes closed condition and to rest quietly with their eyes fixated on a visual fixation point during the eyes open condition. An additional and likely consequential explanation for the differences in results is the disparity in sample sizes between the two studies (see Kharabian Masouleh et al., 2019, for recent work on the inconsistency of replication among studies with differing sample sizes). In any event, the large within-person variation in microstate parameters that we observed in this study implies that future researchers should incorporate longer recordings of resting EEG to obtain sufficiently reliable estimates of microstate dynamics if the goal is to compare microstates between persons. It will also be valuable for future studies to begin to quantify the heterogeneity of microstate temporal parameters across samples, conditions, and analytic and microstate segmentation methodologies.

In contrast to microstate temporal dynamics, the topographic configurations of microstates are strikingly consistent across published reports (Michel and Koenig, 2018). For the present study, microstate clusters A through E matched canonical patterns found in the literature (Michel and Koenig, 2018), including those observed in studies using data-driven approaches to determine an optimal number of clusters (Bréchet et al., 2019; Custo et al., 2017). Together, these five clusters explained 85% of the variance among subject-level cluster centroids, and 63% of the topographic variance (on average) seen in participants’ resting EEG recordings. Nevertheless, a sizeable amount of unexplained topographic variance remained. It is possible that additional microstate configurations, not among the optimal data-driven microstate clusters, could have further explained meaningful patterns in the scalp voltage topography. It is also worth reiterating that identical cluster solutions were found for both the eyes closed and eyes open conditions. Though the temporal dynamics of microstates may be modulated by perceptual state, the present data clearly demonstrate that their topographic configurations are not.

Measuring the functional connectivity of the brain at rest is informative for understanding how cognitive functions change across the lifespan (Campbell and Schacter, 2017). Of the individual difference measures that we examined; the largest effects were obtained for age-related differences in microstate parameters. Increases in microstate duration and decreases in occurrence were observed among older adults for all microstate configurations. These findings support and extend previous findings of age-related differences in microstate duration and occurrence (Koenig et al., 2002; Tomescu et al., 2018). In addition, we found that microstates A and B explained more variance in the EEG time series of older adults, whereas microstates C and E explained less variance, compared to the younger age group. Thus, there appears to be strong evidence for aging-dependent differences in the global temporal properties of EEG brain networks, and the potential for differences in the predominance of particular microstate configurations in older adults.

Younger and older adults also differed on measures of personality, mood, and cognitive function. As a result, we controlled for age group differences when exploring associations between microstate temporal parameters and other individual difference measures. Several significant predictors of microstate dynamics were subsequently identified, and affirmatory evidence provided that many correlations with microstate GEV, duration, and occurrence were also equivalent to zero. However, no significant correlations survived control for multiple comparisons, and so we report and interpret these findings with caution. Additional studies will be needed to confirm the reliability and magnitude of these reported correlations. Nevertheless, given our large sample, we are hopeful that they will remain robust to future replication (cf. Schönbrodt and Perugini, 2013).

Personality describes persistent tendencies that organize individuals’ patterns of thought, emotion, and action over time and across situations. It is thus possible that stable individual differences in personality could be reflected in microstate dynamics present even during periods of quiet rest. This premise was supported by a number of correlations between personality facets and microstates, which ranged from .15 to .21 in magnitude. Facets from the five-factor model of personality tended to be more predictive across parameters than the motivational facets of behavioral inhibition and approach. Of these, agreeableness and conscientiousness had the largest number of correlations of magnitude .15 or higher.

Cognitive performance was also predictive of microstate temporal parameters. Microstate duration was positively associated with psychomotor alertness (i.e., faster reaction times) for configurations A and B, and positively associated with response compatibility (i.e., greater response interference) for configurations A, B, C, and E. In prior work, the neural generators of microstates A and B have been linked to the spontaneous activity of visual and auditory sensory networks, respectively (Bréchet et al., 2019; Britz et al., 2010; Custo et al., 2017). As such, our findings would appear to indicate that the speeded responding and detection of target stimuli is facilitated by individual differences in the tendency for sensory networks to predominate at rest. While fast responding is often advantageous, varying response sets can induce interference-related slowing when the goal is to respond as quickly as possible. This might explain why microstate duration was also correlated with stimulus compatibility effects. In a similar vein, slower reaction times but better response conflict resolution were associated with more frequent occurrences of microstate D. This is in line with studies linking microstate D with activity of a fronto-parietal attentional control network (Britz et al., 2010; Custo et al., 2017), and suggests that this global brain state may be associated with attentional control over automatic response tendencies.

A few more conclusions can be made on the basis of our results. First, individual difference correlations (*r*) of .1 and .2 represent effects between the 25^th^ and 50^th^ percentile of normative effects in psychological research and can be interpreted as small to typical in size, respectively (Gignac and Szodorai, 2016). As such, our results would suggest that microstate dynamics are at best fair predictors of at least some facets of personality and cognitive function across individuals. However, it is also likely that other individual difference measures, not included in this study, may have been stronger predictors of microstate activity, and that different aspects of cognitive function may prove more effective in predicting microstate dynamics. Second, our findings suggest that researchers should take care to match experimental groups on age and gender, and to match conditions on perceptual characteristics when examining microstate temporal dynamics. Indeed, the instructions provided to individuals during eyes closed and eyes open periods may contribute substantially to observed patterns of activity (Milz et al., 2016; Seitzman et al., 2017). Studies should report these instructions in detail, even when participants are simply asked to rest quietly. Finally, there is a rich history of research investigating microstates in populations with neurological or psychiatric dysfunction (e.g., Rieger et al., 2016). It could be the case that microstate temporal parameters are more reliable predictors of pathology than they are of differences among healthy individuals. We hope our findings will nonetheless guide researchers in the selection of measures for future studies in these domains.

Microstates represent one way of parsing spontaneous scalp recorded EEG activity into meaningful functional brain states. Other methods of quantifying oscillatory dynamics or EEG functional connectivity may also prove useful as predictors of individual differences in age, mood, personality, or cognitive function. For example, Mahjoory and colleagues (2019) recently investigated the spectral power and long-range temporal correlations (LRTC) of alpha oscillations in a subset of the present study participants. They found that these oscillatory indices, and their corresponding neural source estimations, were associated with memory span and working memory performance, as calculated from a switch-cost score from the Test of Attentional Performance. Alpha oscillatory power and LRTC quantify similar phenomena to those of microstate dynamics, given that oscillations in the alpha frequency largely determine the periodicity and spatial distribution of microstates in the broadband EEG (Javed et al., 2019; Milz et al., 2017; Wegner et al., 2017). It would be instructive for future studies to more clearly establish functional connections among these various methods. In addition, future studies should continue to investigate the brain generators supporting the waking state under resting task conditions using distributed source estimation techniques. It is our hope that the sharing of large publicly available EEG data sets will spur the advancement of additional analytic methodologies, as has been seen with rapid developments in our understanding of functional and structural fMRI networks resulting from large scale data sharing initiatives.

Despite their topographic similarity across individuals and studies (Michel and Koenig, 2018); their association with fMRI-derived resting state networks (Bréchet et al., 2019; Britz et al., 2010; Custo et al., 2017); and the clear neurophysiological interpretation of microstate temporal parameters (Khanna et al., 2015; Murray et al., 2008)—the functional significance of microstates remains largely unknown in the resting state context. Ultimately, careful phenomenological investigation and experimental manipulation of conscious cognitive acts will be needed to better understand the millisecond fluctuations of microstates and their real-time associations with felt experience and cognition (Varela, 1996). We hope this study will motivate further work on the functional significance of microstates and guide new investigations into the neural correlates of cognition at the millisecond temporal scale.

## Acknowledgements

We thank the Mind-Body-Emotion group at the Max Planck Institute for Human Cognitive and Brain Sciences for collecting and making these data available online. We utilized the freely available Cartool software toolbox (cartoolcommunity.unige.ch) programmed by Denis Brunet, from the Functional Brain Mapping Laboratory, Geneva, Switzerland, and supported by the Center for Biomedical Imaging of Geneva and Lausanne.

## Data Availability Statement

The data accompanying this report are available in the OSF repository and can be found at: https://osf.io/b3uem/

